# *Ex vivo* intestinal permeability assay (X-IPA) for tracking barrier function dynamics

**DOI:** 10.1101/2022.11.08.515584

**Authors:** Hadar Bootz-Maoz, Ariel Simon, Sara Del Mare-Roumani, Yifat Bennet, Danping Zheng, Sivan Amidror, Eran Elinav, Nissan Yissachar

**Affiliations:** The Goodman Faculty of Life Sciences, Bar-Ilan University, Ramat-Gan, 5290002, Israel; Bar-Ilan Institute of Nanotechnology and Advanced Materials, Bar-Ilan University, Ramat-Gan, 5290002, Israel; Systems Immunology Department, Weizmann Institute of Science, 234 Herzl Street, Rehovot, 7610001, Israel; Microbiome & Cancer Division, Deutsches Krebsforschungszentrum (DKFZ), Neuenheimer Feld 280, 69120 Heidelberg, Germany

## Abstract

The intestinal epithelial barrier facilitates homeostatic host-microbiota interactions and immunological tolerance. However, mechanistic dissections of barrier dynamics following luminal stimulation pose a substantial challenge. Here, we describe an *ex-vivo* intestinal permeability assay, X-IPA, for quantitative analysis of gut permeability dynamics at the whole-tissue level. We demonstrate that specific gut microbes and metabolites induce rapid, dose-dependent increases to gut permeability, thus providing a powerful approach for precise investigation of barrier functions.

## Main text

The gut microbiota is confined to the lumen by a single layer of intestinal epithelial cells (IECs). Continuous replenishment of the IEC layer by Lgr5+ intestinal stem cells and physical reinforcement by inter-epithelial tight-junction (TJ) proteins are crucial to maintain barrier integrity and tissue homeostasis^1^. Interference with epithelial barrier integrity (e.g. by pathogenic microbes or drugs) leads to enhanced gut permeability and enables translocation of previously confined (and potentially harmful) microorganisms into the tissue and to distal locations including secondary lymph nodes and liver, where they trigger inflammatory responses^2–4^. Indeed, disruptions to epithelial barrier functions and a ‘leaky gut’ are implicated in both intestinal and extraintestinal pathologies, including allergic and autoimmune conditions, inflammatory bowel diseases, irritable bowel syndrome and cancer^5^.

Existing methods for evaluation of epithelial barrier functions and gut permeability include trans-epithelial electrical resistance (TEER) assay, measurement of ion transport in an Ussing chamber, and oral gavage of animal models with fluorescently labeled molecules followed by quantification of serum fluorescence^6^. However, these approaches do not support real-time perturbations and readouts in an intact, whole-tissue setting, and thus do not provide temporal information of barrier dynamics.

To finely dissect host-microbiota interactions, we developed a unique gut organ culture system that maintains the naïve intestinal tissue architecture and provides tight experimental control^7^. We have demonstrated that this system is ideal for investigating rapid intestinal responses to specific bacterial strains^7^, whole human microbiota samples^8,9^, drugs and metabolites^10^. The emerging realization that the gut microbiota and its associated metabolites regulate intestinal barrier functions^10–12^ drove us to investigate whether the advantages of the gut organ culture system could be harnessed to uncover mechanisms underlying microbial regulation of gut permeability.

Here, we aimed to measure rapid and subtle changes to gut permeability following luminal stimulation, in multiple gut tissues, under tightly controlled experimental conditions and in high temporal resolution. We developed and optimized an *ex vivo* intestinal permeability assay (X-IPA) in which fluorescein isothiocyanate (FITC)-dextran is infused directly into the lumen of the cultured gut tissues (Fig. 1A). Migration of luminal FITC-dextran to the extraintestinal culture medium depends on integrity of the epithelial barrier. Thus, fluorescence intensity of the extraintestinal medium corresponds with epithelial barrier integrity and permeability.

**Figure 1.**
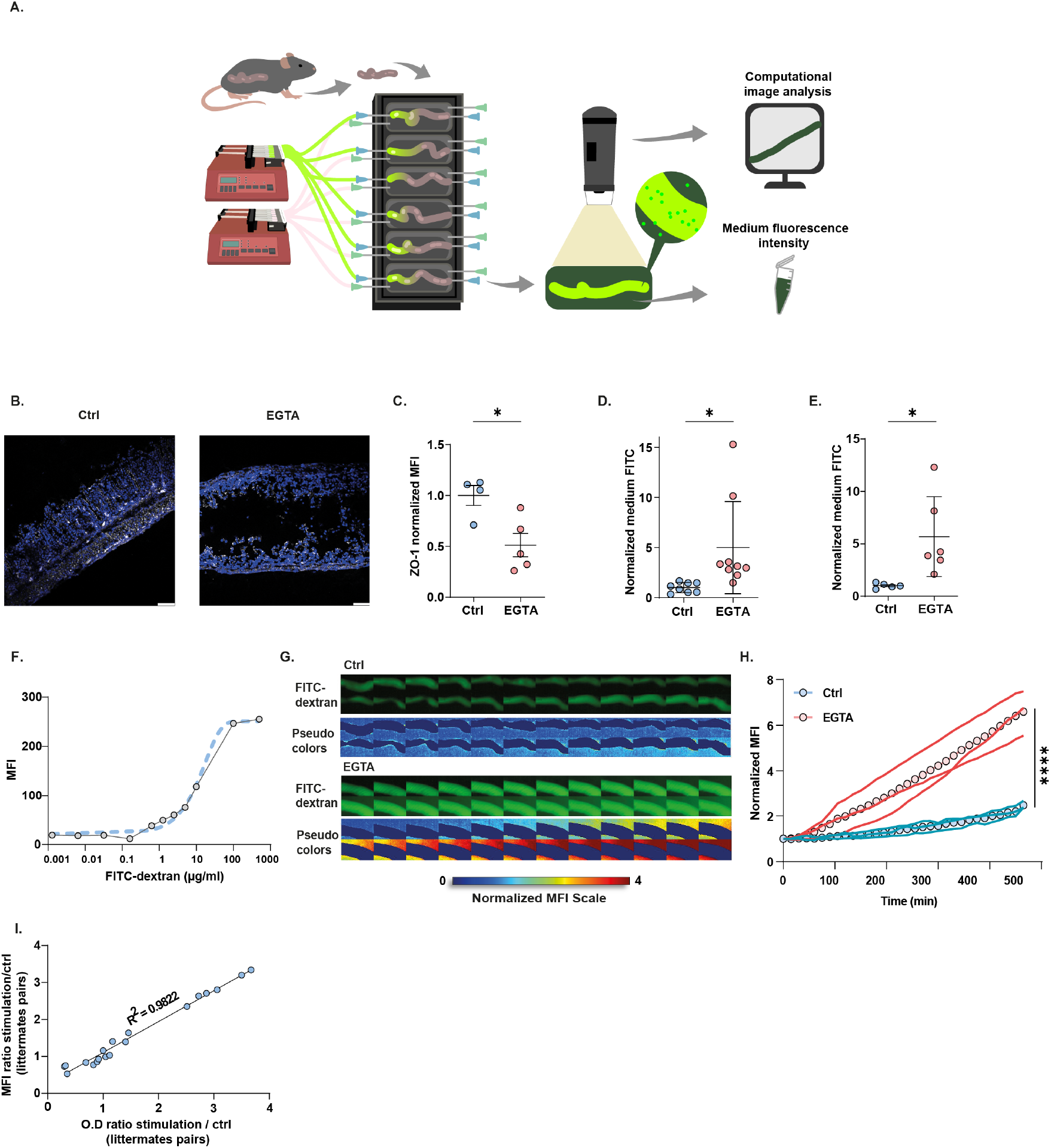
Development of an *ex vivo* intestinal permeability assay (X-IPA). **(A)** Intact intestinal mice tissues are connected to the gut organ culture device and FITC-dextran is infused into the gut lumen. Diffusion over time is quantified by fluorescence intensity of the extra-intestinal medium (time-lapse imaging) and at the experiment end-point (fluorometer), as an indicator of gut permeability degree and temporal dynamics. **(B-C)** Confocal imaging of colon segments cultured for 8h, immuno-stained for ZO-1 (white; DAPI nuclear stain in blue) **(B)** and quantification of ZO-1 MFI following EGTA infusion (p=0.03) **(C)**. **(D-E)** Normalized extraintestinal medium fluorescence of gut cultures infused with EGTA or sterile medium (Ctrl) at 2h (p=0.02) **(D)** and 8h (p=0.04) **(E)**. **(F)** Dynamic range of fluorescence detection by Dino-Lite digital microscope (dashed line - sigmoid nonlinear fit). **(G)** Filmstrips showing gut organ cultures infused with FITC-dextran only (Ctrl) or together with EGTA. The pseudo-color-imaging illustrates MFI quantification of FITC-Dextran concentrations in the extraintestinal medium. Frames are separated by 15 minutes. **(H)** Single colon time traces showing normalized MFI of the extraintestinal medium in tissues infused with EGTA or sterile medium (Ctrl). Circles represent average permeability rates. **(I)** MFI and medium fluorescence intensity (O.D) are tightly correlated (*R*^2^ = 0.9822).

In a set of proof-of-principle experiments, we measured gut permeability *ex vivo* in response to luminal introduction of EGTA, a chemical agent known to disrupt epithelial barrier integrity. Colon tissues were dissected from 14d-old mice reared under specific-pathogen-free (SPF) conditions, and connected to the gut culture device as previously described (six tissues per device; Fig. 1A)^7,8,13^. Culture medium containing FITC-dextran (4kDa) was infused directly into the cultured colons’ lumens either with or without EGTA (25mM). Compared with controls, luminal EGTA infusion disrupted epithelial barrier integrity, as indicated by marked decreases in ZO-1 TJ protein expression levels (Fig. 1B-C). To evaluate whether EGTA-induced barrier disruption affected gut permeability, FITC-dextran fluorescence intensity within the extraintestinal medium was measured following 2h and 8h culture. Luminal infusion of EGTA resulted in a rapid and significant increase in extraintestinal FITC-dextran after only 2h, and continued following 8h of luminal stimulation, compared with controls (Fig. 1D-E), supporting the utility of the X-IPA system in evaluating changes to gut permeability.

A major advantage of the gut organ culture system is the ability to track rapid intestinal responses to experimental perturbations, over time. To quantify barrier dynamics, we added three upright fluorescence digital microscopes (Dino-Lite) to the gut culture system setup, where each microscope tracks two cultured gut fragments (Fig. 1A). Using this set-up, we could efficiently detect subtle changes in medium concentrations of FITC-dextran by quantifying mean fluorescence intensity (MFI) measurements (dynamic range: 1-100 ug/ml; Fig. 1F).

We acquired time-lapse movies of gut organ cultures following luminal introduction of EGTA, at a temporal resolution of 5 min (for 2h experiments) or 15 min (for 8h experiments) (Fig. 1G-H; Supplementary video 1). For automated analysis and quantification, we developed a MATLAB-based image analysis tool and user-friendly interface (X-IPA analyzer; Supplementary Fig. 1 and extended data). This software allowed us to generate dozens of single-tissues time traces and quantify extraintestinal medium MFI, for accurate and comparable characterization of gut permeability dynamics in multiple tissues, over time (Fig. 1G). Automated analysis of time-lapse movies revealed a slow and continuous increase in extraintestinal MFI in non-stimulated tissues (average slope m=0.17), which we interpret as steady-state gut permeability rate under culture conditions. In contrast, EGTA-infused tissues displayed a rapid increase in extraintestinal MFI (average slope m=0.75), observed already at 2h post-stimulation (Fig. 1H; Supplementary video 1). We analyzed FITC-dextran spatial distribution along each intestinal fragment to validate tissue integrity and exclude potential leaking of FITC-dextran from the input or output ports (Supplementary Fig. 2). Further, medium fluorescence at the experiment endpoint (measured by fluorometer) and corresponding image MFI (calculated by image analysis) were well correlated (R^2^=0.9822), demonstrating that time-lapse imaging and analysis can reliably track and quantify gut permeability dynamics (Fig. 1I).

Next, to explore whether the X-IPA system could detect changes to gut permeability induced by intestinal metabolites, gut organ cultures were infused with putrescine, a polyamine compound produced by the gut microbiota which we recently identified as a potent disruptor of the intestinal epithelial barrier^10^. At 4h post-stimulation, we observed a dose-dependent increase in gut permeability in putrescine-infused organ cultures (average slopes: 33mM m≈0; 66mM m=0.28; 100mM m=0.42), compares with controls (Fig. 2A). Automated image analysis revealed a gradual increase in the FITC-dextran diffusion rate (steeper linear slope) in response to increasing putrescine concentrations (Fig. 2B-C; Supplementary video 2). These results could support a model in which TJ opening is a continuous process linearly tuned by luminal signal intensity (putrescine concentration) subsequently resulting in increased permeability.

**Figure 2.**
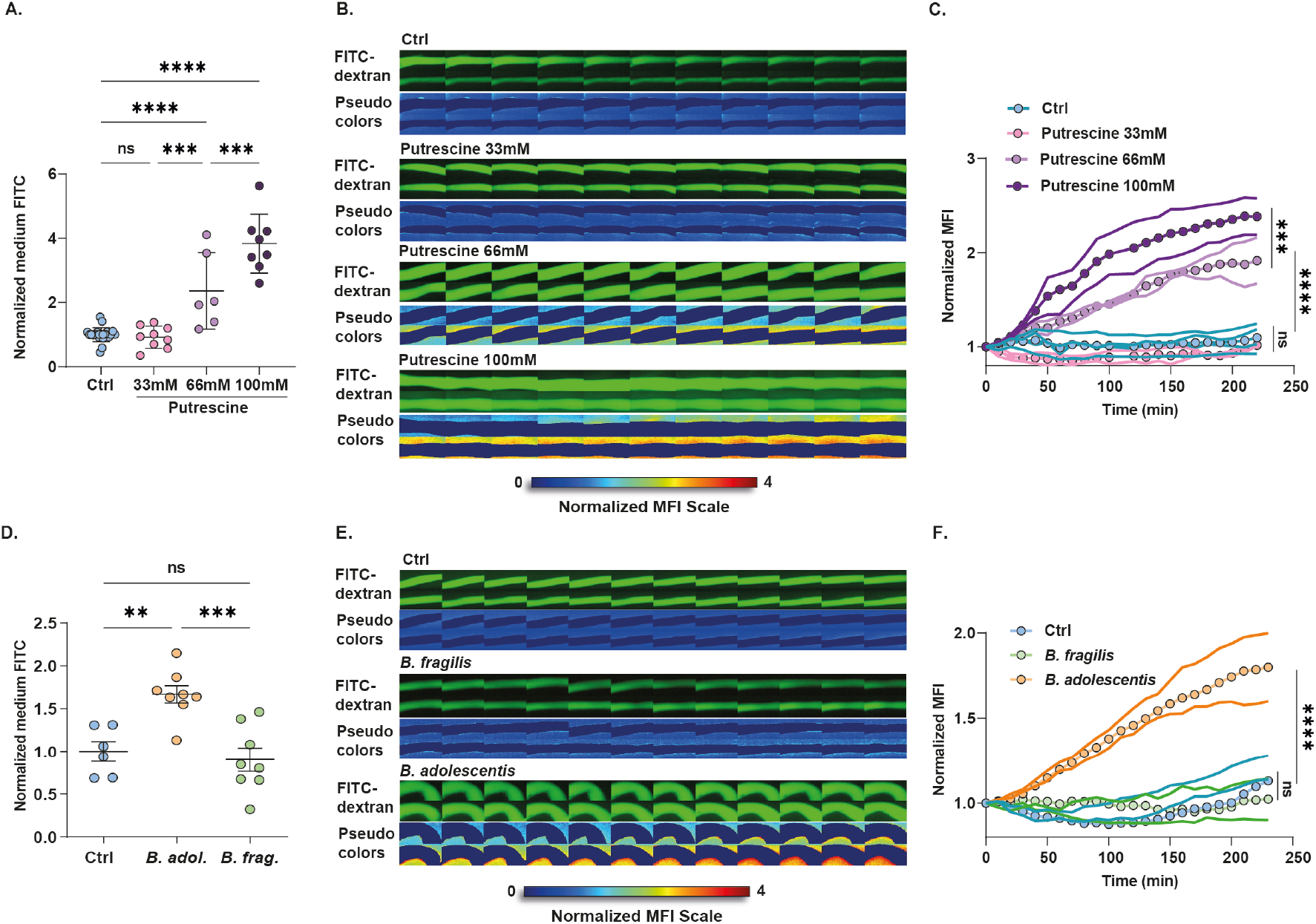
Rapid modulation of gut permeability by metabolites and gut bacteria. **(A, D)** Normalized extraintestinal medium fluorescence of gut cultures infused with increasing putrescine concentrations **(A)** or equivalent load of *B. adolescentis* or *B. fragilis* cultures **(D)**, at 4h post-infusion (normalized to internal control). **(B, E)** Filmstrips showing gut organ cultures infused with FITC-dextran only (Ctrl) or with putrescine **(B)** or microbial cultures **(E)**. The pseudo-color-imaging illustrates MFI quantification of FITC-dextran concentrations in the extraintestinal medium. Frames are separated by 10 minutes. **(C, F)** Single colon time traces showing normalized MFI of the extraintestinal medium in tissues infused with putrescine **(C)** or microbial cultures **(F)** and sterile medium (Ctrl). Statistical significance was determined by one-way ANOVA, p values: **** < 0.0001; *** < 0.001; ** < 0.01; ns - not significant.

Finally, we sought to examine changes to gut permeability induced by the gut microbiota. We analyzed colonic responses to the anaerobic human commensal *Bifidobacterium adolescentis*, a Th17-inducing microbe^14^ which we recently identified as a diet-sensitive pathobiont that alters TJ integrity and disrupts gut barrier functions *in vitro* and *in vivo*^9^. We infused medium containing FITC-dextran and equivalent amounts of *B. adolescentis* or *B. fragilis* (an anaerobic human symbiont that does not induce Th17 development^15^ nor disrupt the epithelial barrier^9^) into the lumen of colonic organ cultures. We observed an increase in extraintestinal MFI in response to barrier-disrupting *B. adolescentis*, but not to *B. fragilis*, at 4h post-stimulation (Fig. 2D). In agreement, time-lapse movies revealed that luminal introduction of *B. adolescentis* rapidly increased extraintestinal MFI (average slope m=0.22), while tissues infused with *B. fragilis* were indistinguishable from controls (average slope m≈0) (Fig. 2E-F; Supplementary video 3).

Taken together, we present a unique organ culture-based gut permeability assay. The X-IPA system provides quantitative insights into the effects of luminal perturbations on gut permeability at the whole-tissue level, thus bridging the gap between simplified *in vitro* assays and complex *in vivo* animal models. Moreover, this system allows, for the first time, to quantify dynamic behaviors of the intestinal barrier as they occur, over time, in high temporal resolution. Additional readouts (including next-generation sequencing, cell sorting, and imaging) applied to the cultured tissues will complement dynamic permeability measurements, thus providing a powerful approach for integrated investigations of host-microbiome interactions. Given the emerging therapeutic potential of barrier modulating agents, we anticipate that the ability to perturb multiple intestinal tissues and to track subsequent gut barrier functions will further advance mechanistic insights and translational discoveries.

## Methods

### Mice

Colons used in the *ex vivo* organ culture system were dissected from sacrificed C57BL/6J (B6) mice. Mice were obtained from Envigo RMS (Israel) and reared in the specific-pathogen-free (SPF) facility at Bar Ilan University (Israel). For gut organ culture permeability assays, colon tissues were dissected from 14-old littermates. All experiments were performed following animal protocols approved by the Bar-Ilan University ethics committee (ethics approval number BIU-BIU-IL-2203-131-1).

### Gut organ culture system and X-IPA permeability assay

Fabrication of the gut organ culture device and gut organ culture experiments were performed as previously described (Yissachar et al., 2017), with the addition of carbon black (Holland-Moran Cas#1333-86-4) into the PDMS (184 ®SYLGARD#761036) mixture, for fabrication of black, opaque gut culture devices. Briefly, intact whole colons were dissected sterilely from 14d-old C57BL/6 mouse littermates reared under SPF conditions. The solid lumen content was gently flushed, and the gut fragment was threaded and fixed over the luminal input and output ports of the gut organ culture device, using sterile surgical thread. The culture device was placed in a custom-made incubator that maintains a temperature of 37°C, and tissue was maintained half-soaked in sterile, serum-free culture medium (Iscove’s Modified Dulbecco’s Medium without phenol-red (IMDM, GIBCO) supplemented with 20% KnockOut serum replacement (GIBCO), 2% B-27 and 1% of N-2 supplements (GIBCO), 1% L-glutamine, 1% non-essential amino acids, 1% HEPES) using a syringe pump. FITC-dextran (4kD) (0.5mg/ml; Sigma-Aldrich Cas#60842-46-8) was resuspended in sterile culture medium without phenol-red and infused into the gut lumen using a syringe pump, with or without EGTA (25mM) (Sigma-Aldrich Cas#13368-13-3), purified bacterial cultures (*B. adolescentis* and *B. fragilis* at 10^7^CFU/ml) or putrescine (Sigma) diluted to final concentrations of 33/66/100mM. Gas outlet in the device lid enabled the flow of a humidified and filtered, medical grade 95% O_2_ / 5% CO_2_ gas mixture into the device. Experiments were terminated at 2h-8h post-stimulation. A fluorescence video microscope (Dino-Lite, Iner Tech Dino-Capture 2.0 AM4115T-GRFBY) acquired time-lapse movies of the cultured tissues and their surrounding culture medium for downstream image analysis and fluorescence quantification. At the experiment end point, the FITC-dextran concentration in the extraintestinal medium was determined by quantifying the fluorescence intensity using a fluorometer (excitation 485nm, emission 520nm). After culture, tissues were subjected to further analysis (including immunofluorescence staining and imaging).

### Image analysis of time-lapse imaging

#### Software development

Automated computerized image analysis of time-lapse movies was performed with the X-IPA Analyzer, a custom written MATLAB software (created in the MathWorks 2020 platform). The main functions in the software are image pre-processing, tissue segmentation and quantification of the green fluorescence intensity of the extraintestinal medium in each chamber, over time (Supp. Figure 1).

#### Experimental design

The experiments were carried out within an organ culture device that contains six chambers, each containing an intact colon tissue and external medium. Three Dino-Lite microscope cameras were placed over the device, such that each one could capture two chambers. The images were taken at regular intervals of 5 min (for 2h experiments) or 15 min (for 8h experiments). A series of images per chamber was analyzed.

#### Analysis algorithm

The algorithm works as follows: The tissue-images are loaded and manually cropped to select pixel-coordinates that represent the measurement area. This area includes the colon-tissue itself and the external tissue-medium. After user-confirmation, the green channel of the RGB model is chosen and the images denoised using a Gaussian filter with a standard deviation of two. The denoised images are segmented by an automated threshold using Otsu’s method from Gray-Level Histograms (Binarization), and the tissue segmentation is expanded by an additional 8 pixels in each direction. Per image, the medium pixels (area outside of the segmentation) are measured, and their mean value is calculated. These mean values are shown in a plot, demonstrating MFI (mean fluorescence intensity) changes over time.

The O.D ratio was calculated by dividing the final concentration of the FITC-dextran (μg/ml) by the final concentration of the sterile medium (Ctrl). The MFI ratio was calculated by dividing the value of the FITC-dextran normalized-MFI final value by the sterile medium (Ctrl) normalized-MFI final value. All steps of analysis can be easily performed with a user-friendly interface (detailed user manual is provided in the online extended data).

### Bacterial cultures

*B. adolescentis* and *B. fragilis* were obtained from DSMZ (Germany), and bacterial classification was further validated by Sanger sequencing of the 16S gene. Microbes were incubated overnight in rich liquid medium (2% proteose peptone (Thermo Fisher Cat#LP0085), 0.5% NaCl (Mercury Cat#1064041) and 0.5% yeast extract (Merck Cas#8013-01-2)) supplemented with 250 mg/ml glucose (Mercury Cas#50-99-7), 250 mg/ml K2HPO4 (Holland Moran Cas#7758-11-4), 50 mg/ml L-cysteine (Merck Cas#52-90-4), 5 mg/ml Hemin (Mercury Cat#3741) and 5ul/ml vitamin *K*_1_ (Merck Cas#84-80-0)) under anaerobic conditions.

### Immunofluorescence staining and confocal imaging

For tissue section staining, intestinal tissues were embedded in OCT and stored at −80°C. The fresh-frozen tissues were sliced using a cryostat (CM1950) into 7 μm sections. The sections were fixed with cold acetone (100%) for 20 minutes, washed twice in PBS and blocked (10% donkey serum, 0.1% triton in PBS) for 1h at room temperature. Tissue sections were then stained for ZO-1 (Invitrogen Cat 40-2200) overnight at 4°C. Excess antibody was washed three times and incubated with secondary antibody (Cy3 anti rabbit Jackson-immuno Cat#711-165-152) for 1h at room temperature. After washing, the sections were stained with DAPI (Merck Cat#1246530100). and mounted (fluoroshield) using an antifade reagent (Sigma F6182).

Tissue sections were visualized using a confocal fluorescence microscope (Leica) and processed and analyzed using ImageJ software. To measure the ZO-1 intensity only in the periphery of the intestinal epithelium, an ImageJ macro was employed, using the following workflow: The images were first Gaussian smoothed (sigma=2), and then segmented by the percentile auto threshold algorithm using the E-cadherin staining. Binary holes were filled to obtain a uniform layer. Next, an ROI (region of interest) band, representing the edge of the intestinal crypt, enclosing the ZO-1 staining, was created as follows: First, the segmented tissue was duplicated and eroded to get a thinner segment. Second, the segmented tissue image was duplicated and dilated to get a wider segment. Finally, the eroded image was subtracted from the dilated image to yield a band, representing the tissue edge. Note, that while the macro allows adjustment of the number of pixels to erode and dilate, once optimal numbers were determined for our images to cover ZO-1, these factors were kept constant for all treatments and for each experiment type. Finally, Analyze Particles was employed to measure the intensity of the total ZO-1 (Cy3) in the ROI.

The intensity of ZO-1 in cultured cells was measured as follows: Smoothing on the ZO-1 image (channel 3 Cy3) using Gaussian blur (sigma=1) was applied. Next, the background was subtracted (rolling ball=10). The cells were then segmented using the automatic Otsu algorithm. Finally, the intensity of the ZO-1 marker was calculated according to the binary segmented image.

## Supporting information

X-IPA analyzer user manual

X-IPA analyzer.exe

---

**Supplementary figure 1:**
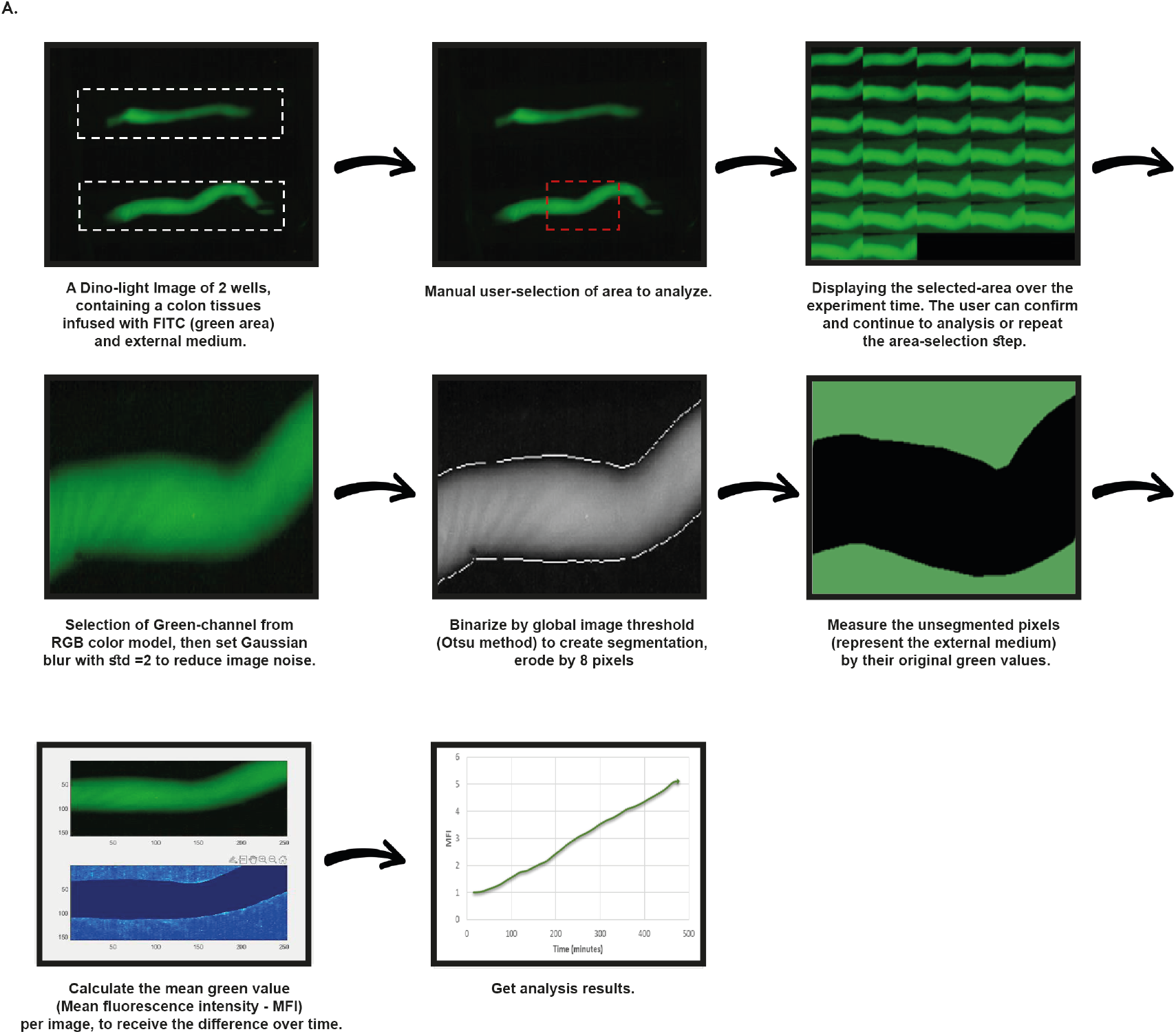
Schematic representation of automated image analysis procedure using the X-IPA analyzer. The tissue-images are loaded and cropped to mark the measurement-area. These coordinates include the intestinal tissue and the external surrounding medium. After user confirmation, the green channel of the RGB model is chosen and the images denoised using Gaussian filter, with a standard deviation of two. The denoised images are segmented by an automated threshold using Otsu’s method from Gray-Level Histograms (Binarization), and the tissue segmentation is expanded by extra 8 pixels. Per image, the medium pixels (area outside of the segmentation) are measured, and their mean fluorescence value is calculated. These mean fluorescence values are shown in a plot, demonstrating the MFI (mean fluorescence intensity) changes over time.

**Supplementary figure 2:**
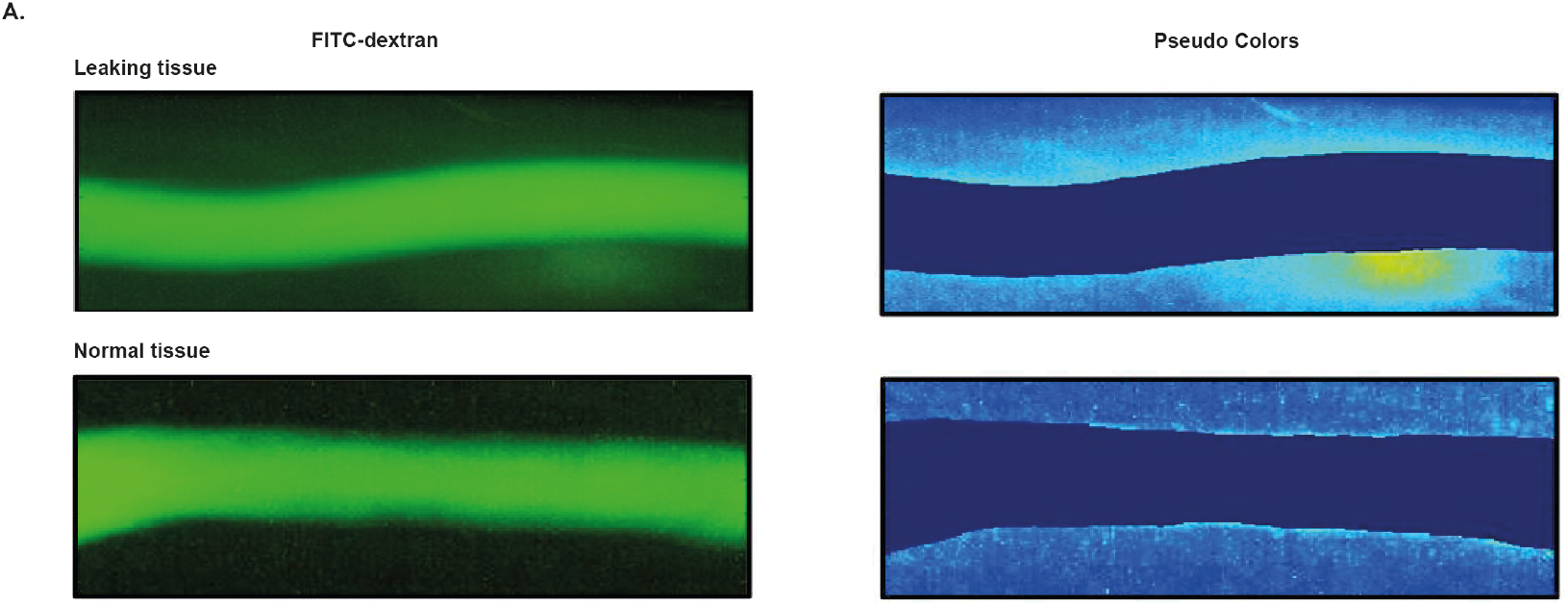
Quality control – detection of abnormal extraintestinal fluorescence. The X-IPA analyzer software allows the user to identify abnormal MFI values, which may result from tissue damage during surgery and not from increased intestinal permeability. The interactive pseudo-colors allow the user to detect the leakage of FITC-dextran from the tissue throughout the time lapse movie. As an example, the upper images show a damaged, leaky tissue. The leaky area appears stronger according to the pseudo-colors scale, and concentrated within a defined region of the extraintestinal medium. The bottom images show a normal, intact tissue, that is surrounded by medium with normal fluorescence distribution.

**Supplementary video 1: EGTA rapidly increases gut permeability**

A time lapse movie showing colon tissues infused with FITC-dextran for 8 hours, reveals a dramatic increase of fluorescence in the medium of the EGTA infused tissues compared to Ctrl (sterile medium only). Here, 3 EGTA-infused tissues and 3 medium-infused tissues were measured. The FITC-dextran animation shows the increasing of fluorescence in a representative tissue, as captured by the Dino-light microscope lens over time. The Pseudo-colors animations shows the increasing of fluorescence in the tissue, as it looks in the X-IPA analyzer software over time. Frames are separated by 15 minutes.

**Supplementary video 2: Putrescine rapidly increases gut permeability, in a dose-dependant manner**

A time lapse movie showing colon tissues infused with FITC-dextran for 4 hours, reveals an increase in extraintestinal medium fluorescence following infusion of putrescine compared to Ctrl (sterile medium only). The FITC-dextran animation shows the increasing of fluorescence in a representative tissue, as captured by the Dino-light microscope lens over time. The Pseudo-colors animations shows the increasing of fluorescence in the tissue, as it looks in the X-IPA analyzer software over time. Frames are separated by 5 minutes.

**Supplementary video 3: *B. adolescentis*, but not *B. fragilis*, rapidly increases gut permeability**

A time lapse movie showing colon tissues infused with FITC-dextran for 4 hours, reveals an increase in extraintestinal medium fluorescence following infusion of *B. adolescentis* compared with tissues infused with *B. fragilis* or Ctrl (sterile medium only). The FITC-dextran animation shows the increasing of fluorescence in a representative tissue, as captured by the Dino-light microscope lens over time. The Pseudo-colors animations shows the increasing of fluorescence in the tissue, as it looks in the X-IPA analyzer software over time. Frames are separated by 5 minutes.

## Acknowledgements

This work is dedicated to the loving memory of Prof. Nir Friedman z”l - a dear friend, a mentor and a source of inspiration. We thank Sondra Turjeman for critical editing of the manuscript, Hagar Morad, Shir Ben-Aaron and Nadav Vernia for computational image analysis and members of the Yissachar lab for insightful discussions. This work was supported by the Israel Science Foundation (grant No.1384/18), the Israel Science Foundation-Broad Institute Joint Program (grant No. 2615/18), and the Gassner Fund for Medical Research, Israel.

## References

1. Turner, J. R. Intestinal mucosal barrier function in health and disease. Nat. Rev. Immunol. 9, 799–809 (2009).

2. Yang, Y. et al. Within-host evolution of a gut pathobiont facilitates liver translocation. Nature 607, 563–570 (2022).

3. Manfredo Vieira, S. et al. Translocation of a gut pathobiont drives autoimmunity in mice and humans. Science 359, 1156–1161 (2018).

4. Ruff, W. E., Greiling, T. M. & Kriegel, M. A. Host-microbiota interactions in immune-mediated diseases. Nat. Rev. Microbiol. 18, 521–538 (2020).

5. Akdis, C. A. Does the epithelial barrier hypothesis explain the increase in allergy, autoimmunity and other chronic conditions? Nat. Rev. Immunol. 21, 739–751 (2021).

6. Schoultz, I. & Keita, Å. V. The Intestinal Barrier and Current Techniques for the Assessment of Gut Permeability. Cells 9, 1909 (2020).

7. Yissachar, N. et al. An Intestinal Organ Culture System Uncovers a Role for the Nervous System in Microbe-Immune Crosstalk. Cell 168, 1135–1148.e12 (2017).

8. Duscha, A. et al. Propionic Acid Shapes the Multiple Sclerosis Disease Course by an Immunomodulatory Mechanism. Cell 180, 1067–1080.e16 (2020).

9. Pearl, A. et al. Diet-induced modifications to human microbiome reshape colonic homeostasis in irritable bowel syndrome. bioRxiv (2021).

10. Grosheva, I. et al. High-Throughput Screen Identifies Host and Microbiota Regulators of Intestinal Barrier Function. Gastroenterology (2020). doi:: 10.1053/j.gastro.2020.07.003

11. Heppert, J. K. et al. Transcriptional programmes underlying cellular identity and microbial responsiveness in the intestinal epithelium. Nat. Rev. Gastroenterol. Hepatol. 18, 7–23 (2021).

12. Tilg, H., Zmora, N., Adolph, T. E. & Elinav, E. The intestinal microbiota fuelling metabolic inflammation. Nat. Rev. Immunol. (2019). doi:: 10.1038/s41577-019-0198-4

13. Azriel, S., Bootz, H., Shemesh, A., Amidror, S. & Yissachar, N. An Intestinal Gut Organ Culture System for Analyzing Host-Microbiota Interactions. J. Vis. Exp. (2021). doi:: 10.3791/62779

14. Tan, T. G. et al. Identifying species of symbiont bacteria from the human gut that, alone, can induce intestinal Th17 cells in mice. Proc. Natl. Acad. Sci. 113, E8141–E8150 (2016).

15. Mazmanian, S. K., Round, J. L. & Kasper, D. L. A microbial symbiosis factor prevents intestinal inflammatory disease. Nature 453, 620–5 (2008).

