## Supplementary figures and images for "*Ex vivo* intestinal permeability assay (X-IPA) for tracking barrier function dynamics"

### Supplementary fig 1

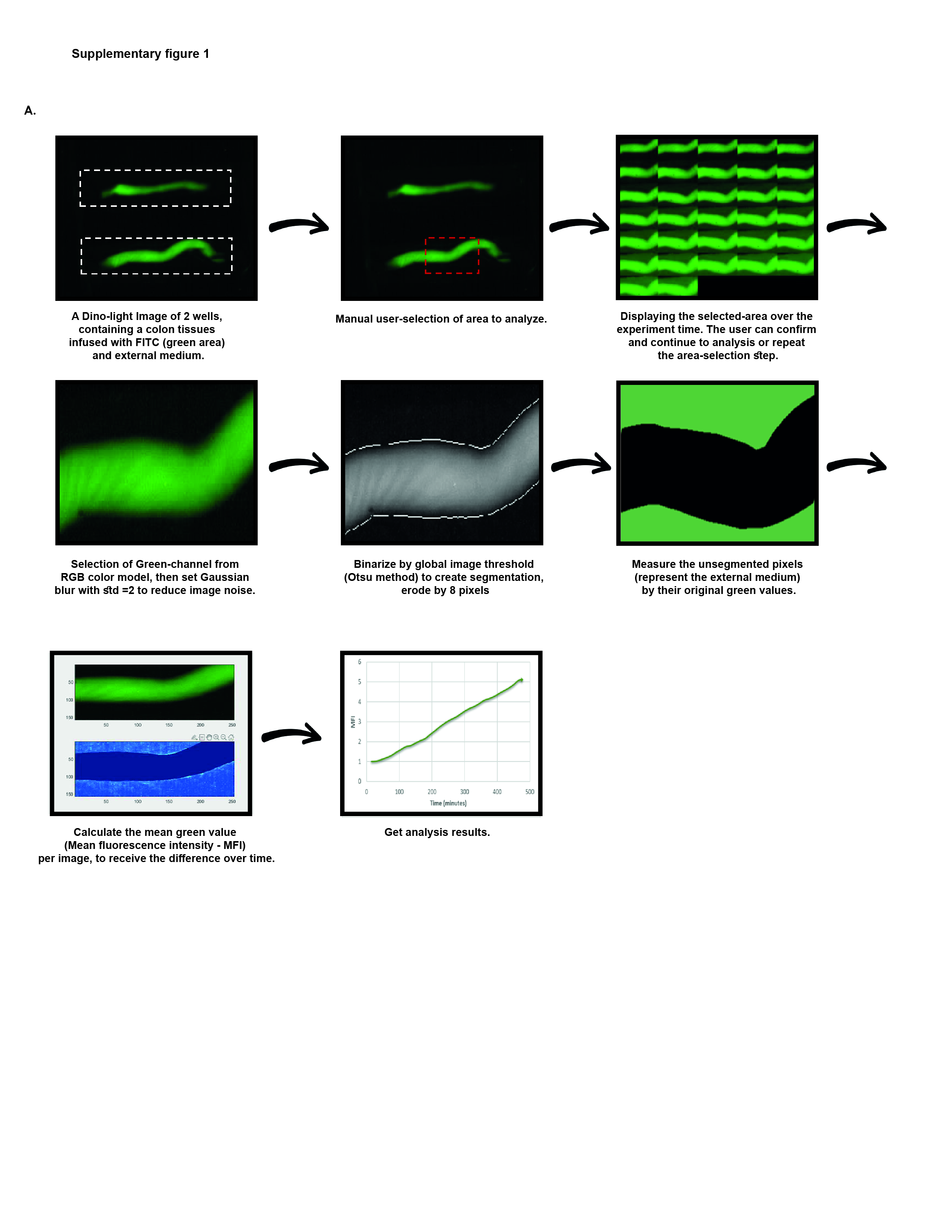

### Supplementary fig 2

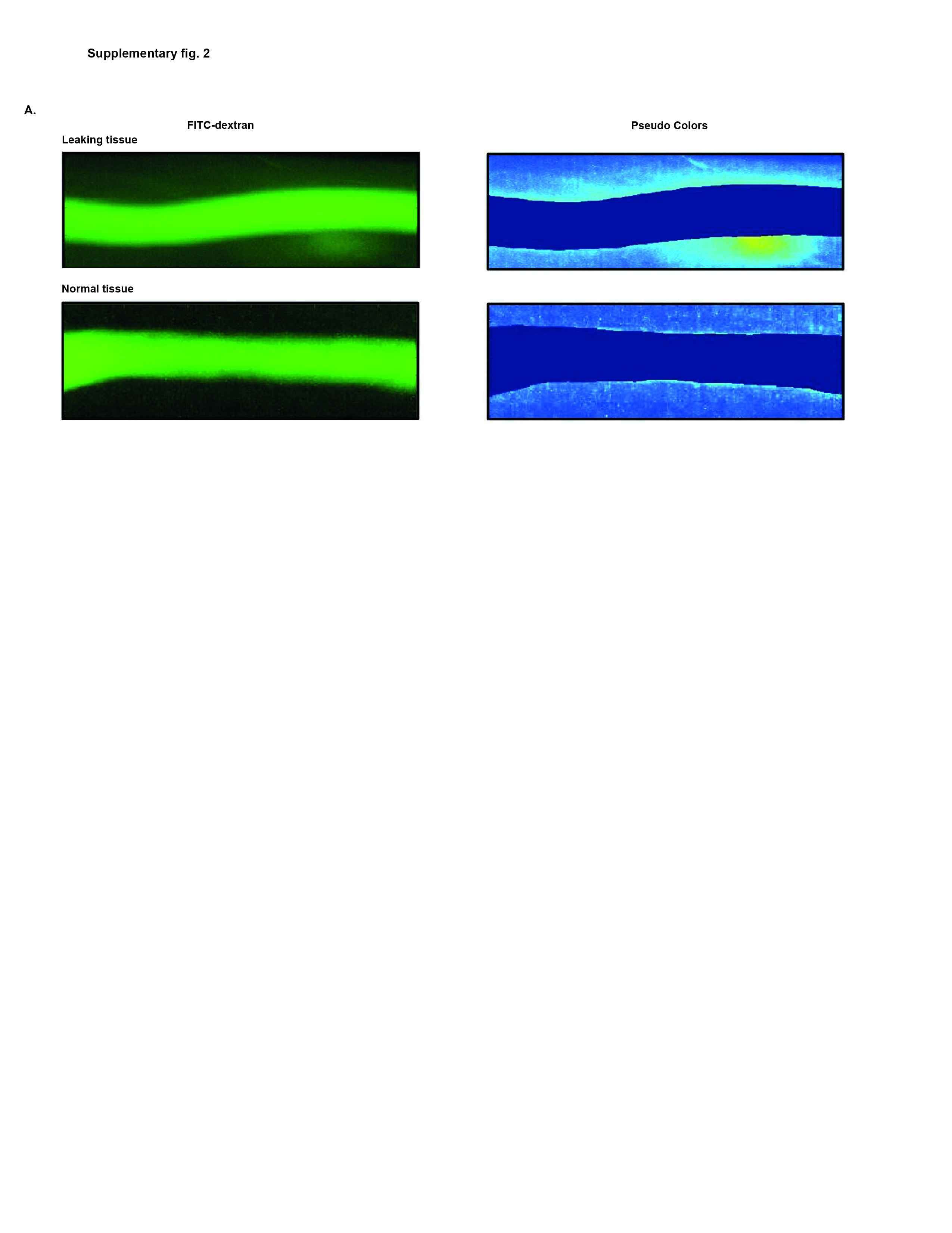
