## Supplementary material for "*Ex vivo* intestinal permeability assay (X-IPA) for tracking barrier function dynamics": X-IPA analyzer user manual

### X-IPA Analyzer – User manual for image analysis

#### Requirements:

- Install the extension "matlab runtime compiler."
  [MATLAB Runtime - MATLAB Compiler - MATLAB (mathworks.com)]
- Before running the program, the images obtained from the "Dino- Lite" microscope must be split into separate folders.
- Install XIPA.exe file on user's computer.

#### Instructions:

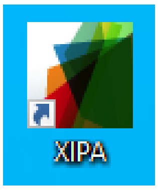


Double-click on the application icon. Wait few seconds until this screen appears:

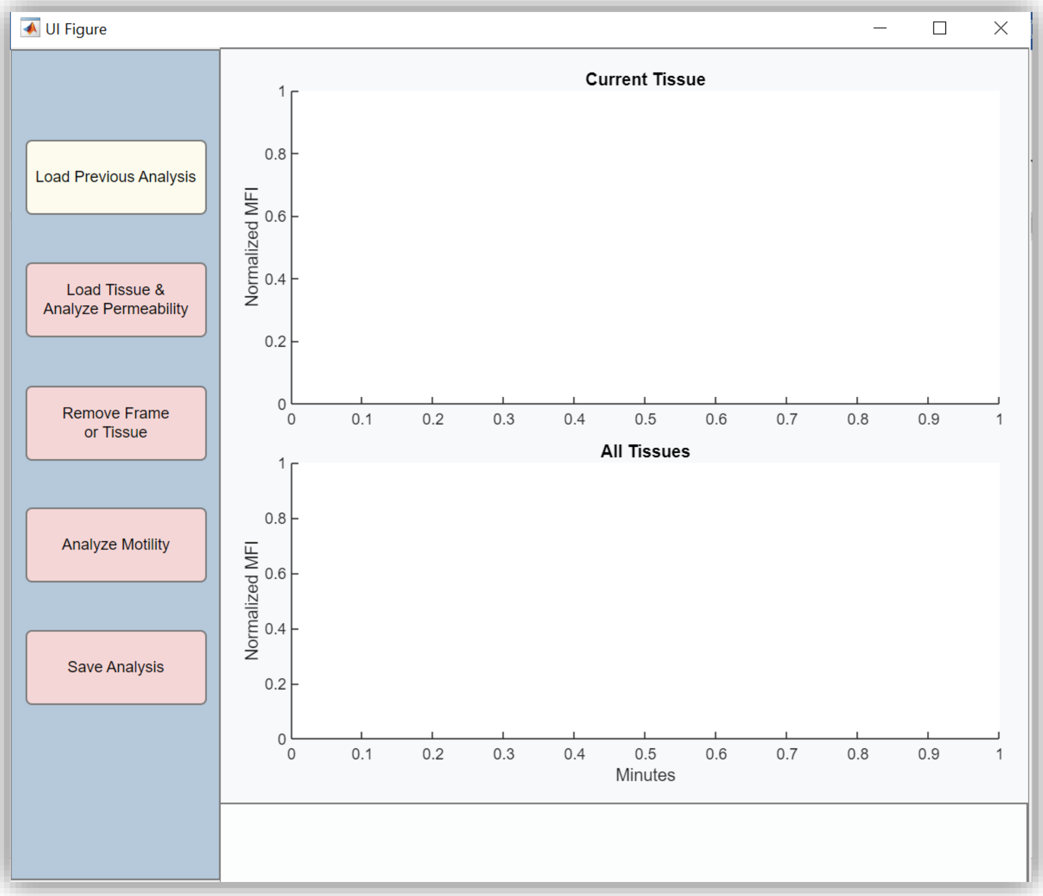


User will be prompted with a window to select the location where they would like to save the results from the analysis.

**Load Tissue & Analyze Permeability button**

Clicking this button will ask the user to select the file where the images are located. After selecting the file, the system will display the following box with the fields that must be filled in for the analysis documentation:

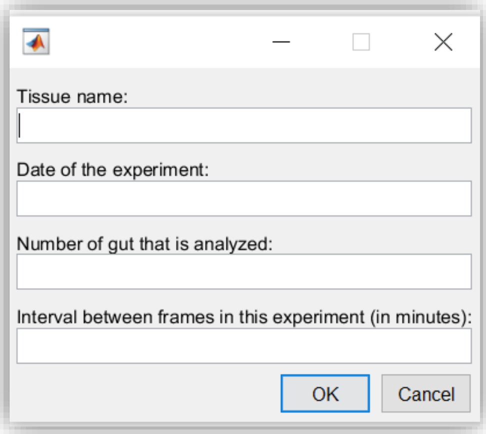


- The name of the tissue loaded into the program (for example- "EGTA")
- The date of the experiment
- The tissue number
- Interval (the time in minutes) between the images in the experiment

If the images are loaded from the Dropbox folder, it may take a few seconds to upload the images.

The program will then upload the series of images and present the first image to the user. The user must select the relevant area of the image for analysis by surrounding it with a red rectangle. Choose a clean and clear area without shadow or the needles that fix the tissue. Make sure that the selected area contains both the relevant intestinal tissue as well as the surrounding medium.

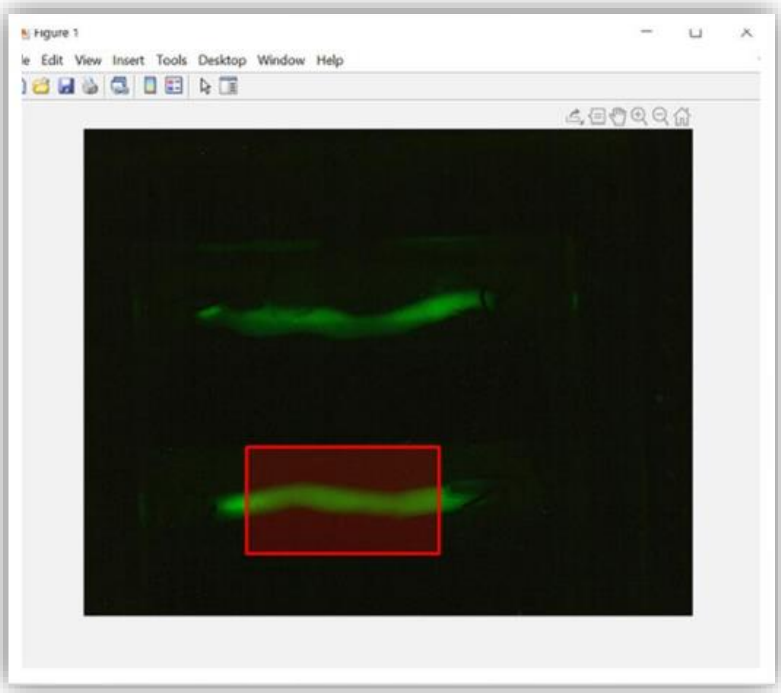


The program will crop the entire series of images according to the coordinates of the rectangle and display them to the user. The user is prompted to decide whether to continue with the current crop, or to re-crop (if the current crop is not optimal).

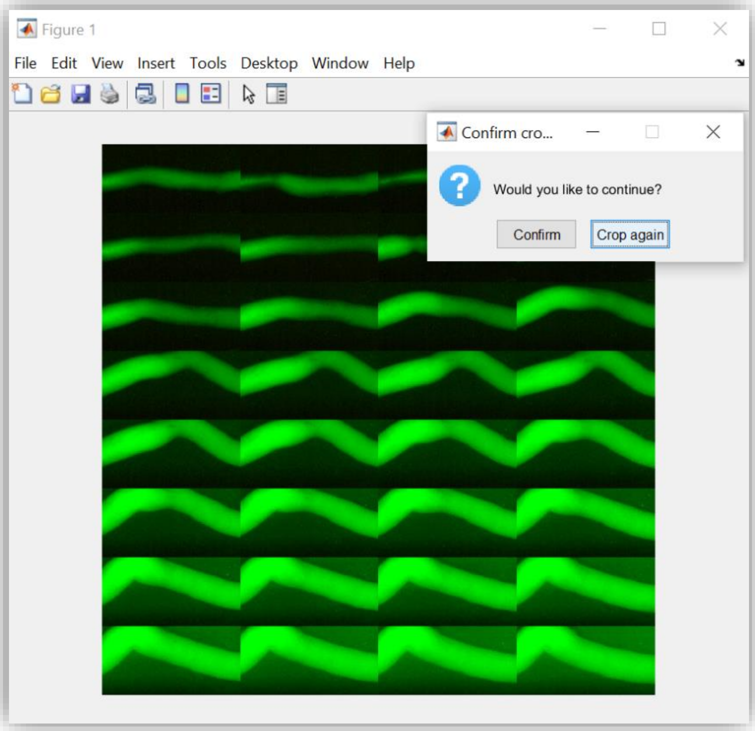


In fact, the user can repeat the cropping process indefinitely by clicking "Crop again." If the user is satisfied with the current images, they may click "Confirm" and the program will proceed to process the images.

During the processing, a video is displayed that simulates the change in the fluorescent intensity in the wall's medium throughout the experiment, as well as a "montage" of the segmentation of all the images. These two outputs are for monitoring the continuation of the analysis.

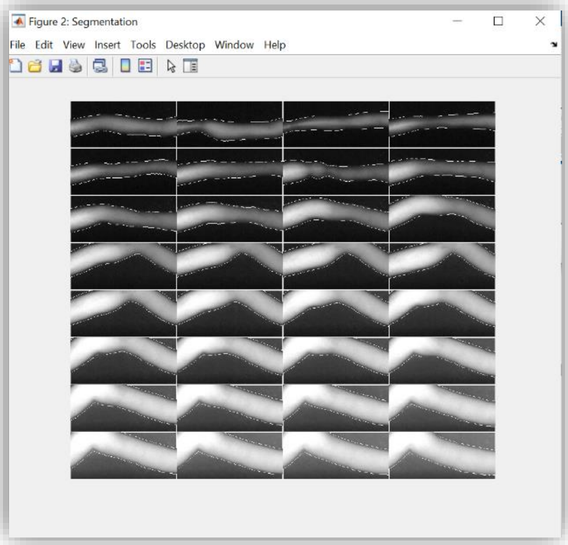


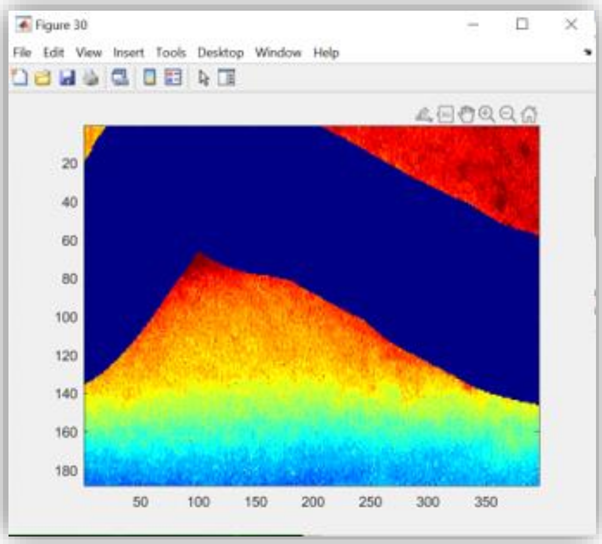


At the end of the video, the user will be notified "analysis done" and a MFI graph will be displayed in the upper panel, depicting the MFI values along the images in the current series.

In the lower panel, the graph of all tissues that have been loaded for analysis will appear. The x-axis values represent the time in minutes, corresponding to the intervals entered by the user when loading the tissue.

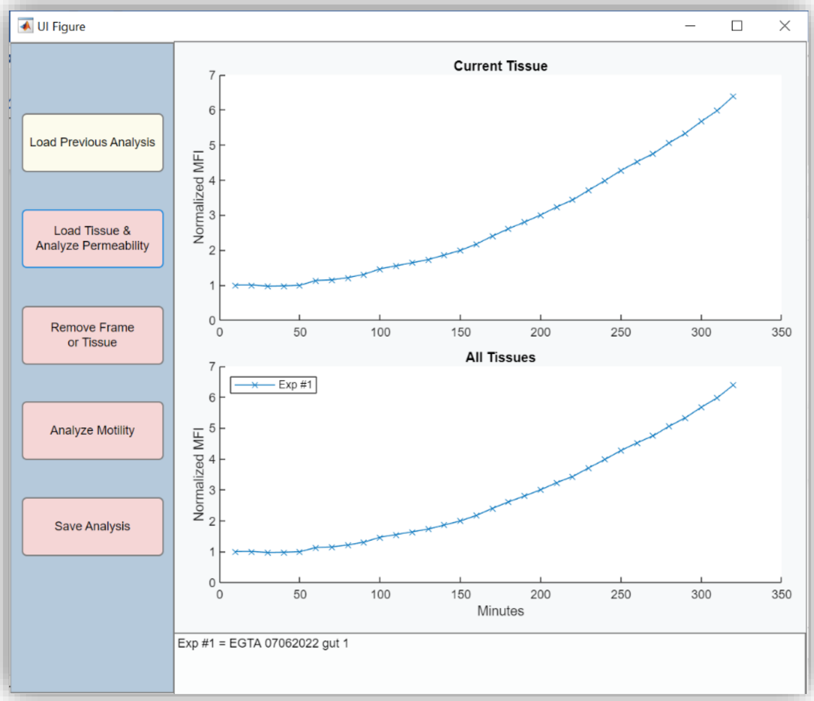


The above steps can now be repeated for as many tissue images as the user wishes to load and analyze together. At the end of the run, graphs will appear describing the MFI values in the medium in each of the tissues.

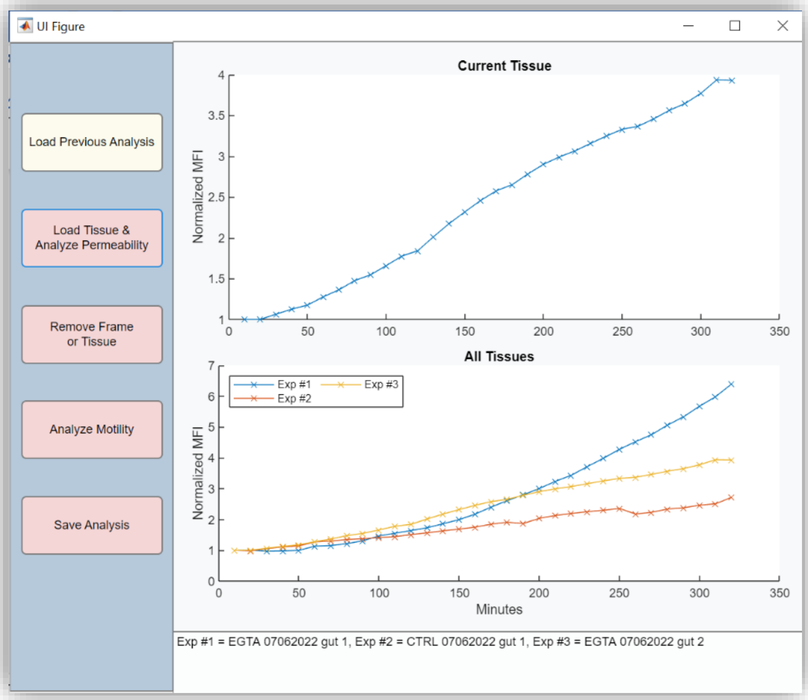

#### Remove Frame or Tissue button

This button can delete an entire tissue from the analysis or conduct a pointwise removal of any point in the graph of one of the experiments. Depending on the need, select the "Frame" button or the "Tissue" button.

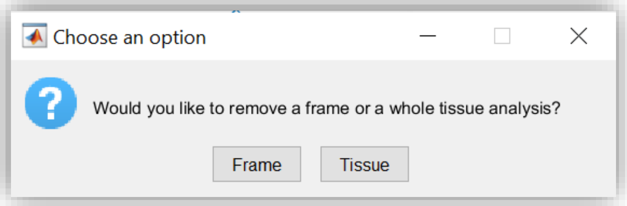


**Tissue deletion:**

When choosing to delete a tissue, a panel is displayed in which the user must enter the number of the tissue they wish to delete (according to the legend that appears in the graph panel of all tissues):

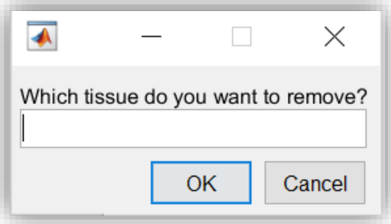


After clicking "OK," the selected tissue is deleted from all graphic panels and its values are removed from the analysis data completely.

**Frame deletion:**

When choosing to remove a point, the user must input the tissue number and the x value (number of minutes) of the point to be deleted. The value of the desired point can be easily obtained by clicking on it in the graph.

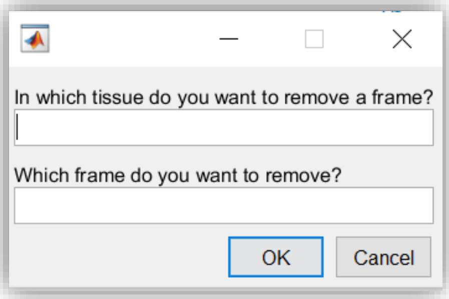


The removal of the point is done by replacing its original value with the average of the two adjacent points. After the selection, the point is "smoothed" both in the graph and in the calculated experimental values.

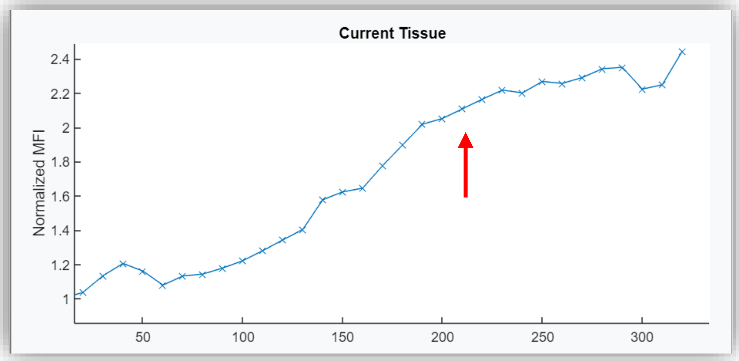

*After removing the Y value of "210" time point*

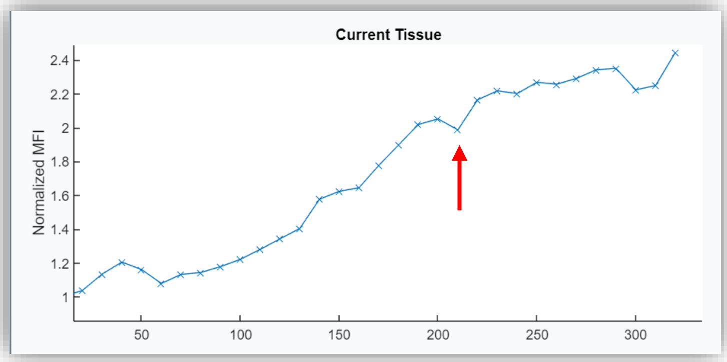

*Before removing the Y value of "210" time point*

- Deleting the first image in the series is not possible since the rest of the graph values are normalized based on it.
- Deleting the last point in the series will lead to lowering the number of time points in the series.

**<u>Load Previous Analysis button</u>**

This button makes it possible to upload images and videos from previous analyses performed by the software into the current analysis data of calculated MFI values. Selecting this button displays a window for choosing the folder containing the Excel file of the data of the previous analysis.

After the selection, the graphs representing the loaded tissues are displayed. At the same time, the segmented images, videos, and MFI data from the loaded analysis are copied to the current results folder.

- The user should pay attention to the value of the intervals of the tissues that are loaded into the system as they may differ from the intervals of the tissues that have previously been loaded.

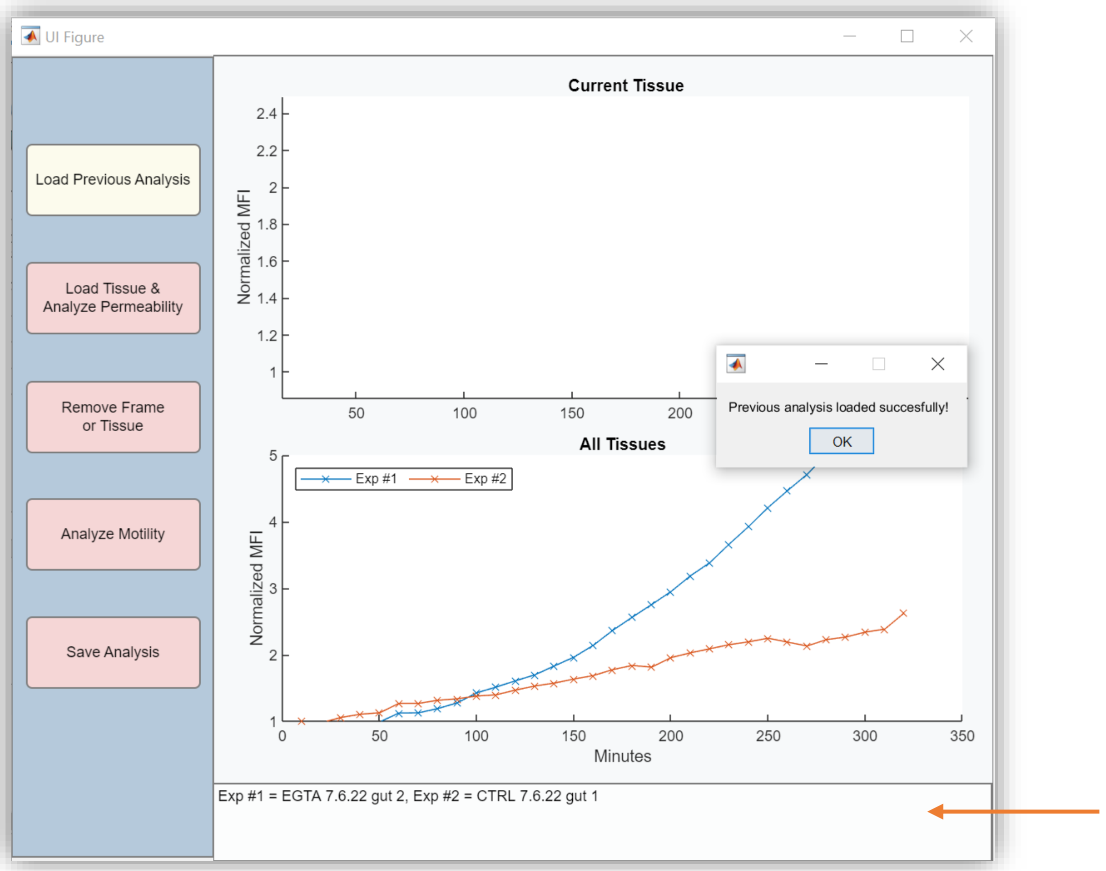


**Please note** - while the program is running, a text window will appear at the bottom of the screen indicating the number of the tissue being loaded, as well as the name of the tissue provided by the user (marked with an orange arrow). Thus, when loading new tissues, previous analyses or removing experiments from the graph, the user can follow and adjust between a curve and a tissue.

#### Save Analysis button

This button enables the saving of the analysis data performed in the application panel.

By clicking the button, a documentation box opens, prompting the user to enter data about the analysis performed before saving:

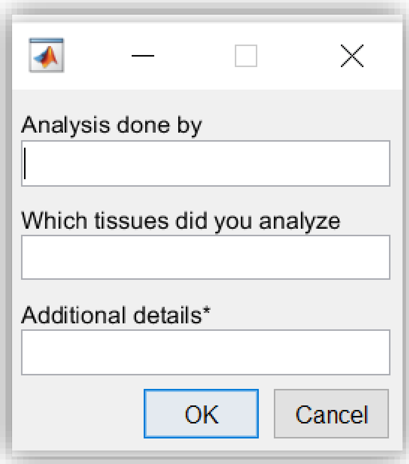

*The "Additional details" field is optional.

The information entered in this box is saved as a text file named "details.txt" in the results folder of the experiment. Now, the program will rename the results folder to the string of tissue names entered by the user. **For example**:

| Name | Date modified |
|---|---|
| ctrl 070622 gut 1 + EGTA 070622 gut 2 + b.adol 070622 gut 1 + b.frag 070622 gut 2 | 6/7/2022 1:25 PM |

The user can change this name once the process is completed.

The results folder contains the series of images per experiment after segmentation, the video simulators and the Excel file named "data MFI". For example:

| Name | Date modified | Type | Size |
|---|---|---|---|
| b.adol 070622 gut 1 animation | 6/7/2022 1:05 PM | AVI File | 903 KB |
| b.adol 070622 gut 1 segmentation | 6/7/2022 1:05 PM | PNG File | 867 KB |
| b.frag 070622 gut 2 animation | 6/7/2022 1:06 PM | AVI File | 959 KB |
| b.frag 070622 gut 2 segmentation | 6/7/2022 1:06 PM | PNG File | 784 KB |
| ctrl 070622 gut 1 animation | 6/7/2022 1:03 PM | AVI File | 1,279 KB |
| ctrl 070622 gut 1 segmentation | 6/7/2022 1:03 PM | PNG File | 734 KB |
| data_MFI | 6/7/2022 1:10 PM | גליון עבודה של Mic... | 7 KB |
| details | 6/7/2022 1:11 PM | Text Document | 1 KB |
| EGTA 070622 gut 2 animation | 6/7/2022 1:04 PM | AVI File | 863 KB |
| EGTA 070622 gut 2 segmentation | 6/7/2022 1:04 PM | PNG File | 783 KB |

The data MFI file contains two sheets:

**"Normalized data" sheet**

Each column contains the name of the tissue given by the user, the value of the intervals entered, and the normalized data for the first frame calculated for the tissue during the current program run (unless removed by the user). It should be noted that in this tab excess frames are "cut", that is, if an experiment with x images and an experiment with x+2 images are analyzed, the program will only keep the values of the x frames to maintain normal matrix dimensions.

**"Raw data" sheet**

This data is not normalized and contains all the frames from each experiment. To maintain the integrity of MATLAB programming, the data appears in rows (can be turned on manually by transpose).

At the end of saving the analysis, a window will pop up indicating as such.

**If the analysis stops before saving the results for any reason, the user can find the calculated results under a folder named yyyy-mm-dd Analysis Result, the date being the day of user access to the software.**
